# An infralimbic cortex neuronal ensemble encoded during learning attenuates fear generalization expression

**DOI:** 10.1101/2024.08.18.608308

**Authors:** Rajani Subramanian, Avery Bauman, Olivia Carpenter, Chris Cho, Gabrielle Coste, Ahona Dam, Kasey Drake, Sara Ehnstrom, Naomi Fitzgerald, Abigail Jenkins, Hannah Koolpe, Runqi Liu, Tamar Paserman, David Petersen, Diego Scala Chavez, Stefano Rozental, Hannah Thompson, Tyler Tsukuda, Sasha Zweig, Megan Gall, Bojana Zupan, Hadley Bergstrom

## Abstract

Generalization allows previous experience to adaptively guide behavior when conditions change. The infralimbic (IL) subregion of the ventral medial prefrontal cortex plays a known role in generalization processes, although mechanisms remain unclear. A basic physical unit of memory storage and expression in the brain are sparse, distributed groups of neurons known as ensembles (i.e., the engram). Here, we set out to determine whether neuronal ensembles established in the IL during learning contribute to generalized responses. Generalization was tested in male and female mice by presenting a novel, ambiguous, tone generalization stimulus following Pavlovian defensive (fear) conditioning. The first experiment was designed to test a role for IL in generalization using chemogenetic manipulations. Results show IL regulates defensive behavior in response to ambiguous stimuli. IL silencing led to a switch in defensive state, from vigilant scanning to generalized freezing, while IL stimulation reduced freezing in favor of scanning. Leveraging activity-dependent “tagging” technology (ArcCreER^T2^ x eYFP system), a neuronal ensemble, preferentially located in IL Layer 2/3, was associated with the generalization stimulus. Remarkably, in the identical discrete location, fewer reactivated neurons were associated with the generalization stimulus at the remote timepoint (30 days) following learning. When an IL neuronal ensemble established during learning was selectively chemogenetically silenced, generalization increased. Conversely, IL neuronal ensemble stimulation reduced generalization. Overall, these data identify a crucial role for IL in suppressing generalized responses. Further, an IL neuronal ensemble, formed during learning, functions to later attenuate the expression of generalization in the presence of ambiguous threat stimuli.

## INTRODUCTION

When confronted with a clear and present threat, organisms must respond appropriately for defense. Sometimes threats are unclear, or ambiguous, which might call for a more generalized response, based on past experience (Richards & Frankland, 2017). In the field of psychology, generalization refers to the transfer of conditioned responses to stimuli that are similar, but not identical, to the original conditioned stimulus (Guttman & Kalish, 1956). Generalization was first described over a century ago (Pavlov, 1927; Watson & Rayner, 1920), is highly conserved across species (Dymond et al., 2015; Orr & Lukowiak, 2008), and has even been proposed as one of the only universal laws in the field of psychology (Shepard, 1987). While generalization is evolutionarily adaptive, overgeneralization of defensive behavior is maladaptive, and relevant to understanding posttraumatic stress disorder (PTSD) and anxiety-related disorders (Cooper et al., 2022; Dunsmoor & Paz, 2015). The fundamental nature of stimulus generalization, in both the study of intrinsic memory processes and clinical disorders, makes the investigation of generalization a central topic in the field of neuroscience (Sangha et al., 2020).

The ventral medial prefrontal cortex (vmPFC), and infralimbic (IL) subregion (cingulate cortex Area 25), is an identified hub in the defensive conditioning circuit (Sotres-Bayon et al., 2006). Functionally, a role for IL in extinction processes is canonical (Giustino & Maren, 2015). A growing number of studies have also indicated IL functionality in generalization (Bayer & Bertoglio, 2020; Corches et al., 2019; Day et al., 2020; Kreutzmann et al., 2020; Kreutzmann & Fendt, 2020; Ng et al., 2023; Sangha et al., 2014; Scarlata et al., 2019). The consensus from these studies is that IL functionality mediates memory specificity by attenuating generalization.

The term neural ensemble or “engram” refers to a sparse, distributed, group of neurons that forms a physical substrate of memory in the brain (Josselyn et al., 2015). An approach for studying neuronal ensembles is the use of molecular genetic tools for the selective expression (or “tagging”) of fluorescent molecules, or optogenetic and chemogenetic actuators, during learning for later visualization and manipulation during memory retrieval (DeNardo & Luo, 2017). One such system is the ArcCreER^T2^ x eYFP mouse line. In this system, activity-regulated cytoskeletal-associated (Arc) gene transcription, a key component in learning and memory-induced synaptic plasticity, leads to the expression of a CreER^T2^ fusion protein which, upon activation with a synthetic estrogen receptor (ER) agonist (i.e., 4-OHT), leads to Cre-induced recombination and expression of a fluorescent molecular tag (Denny et al., 2014).

Generalization has long been proposed to be established during learning (Hull, 1943), although this hypothesis has not been directly tested. Here, we designed a set of experiments to determine whether an IL neuronal ensemble encoding an inhibitory process is established during Pavlovian defensive conditioning. We hypothesize that after learning, this ensemble contributes to memory specificity by reducing conditioned responses to generalization stimuli. The ArcCreER^T2^ x eYFP transgenic mouse system is particularly advantageous for addressing this question because neurons activated during learning can be later manipulated under conditions that promote generalization.

To first establish a role for IL in cued fear memory generalization we used a Designer Receptors Exclusively Activated by Designer Drugs (DREADD) system and CaMKII promoter to bidirectionally control IL excitability during the expression of generalization. In the second set of experiments, we took advantage of the ArcCreER^T2^ x eYFP system to visualize and measure IL neuronal ensembles associated with generalization at recent and remote timepoints following learning. In the final set of experiments, a double-floxed inverse open reading frame (DIO)-DREADD system and hSyn promoter in combination with ArcCreER^T2^ x eYFP transgenic mice was used to transfect DREADDs constructs in Arc-expressing neurons during learning for later synthetic chemical manipulation during memory retrieval.

## MATERIALS and METHODS

### Animals

Adult male and female ArcCreER^T2^ x eYFP mice were used in all experiments. B6.Cg-Tg(Arc-cre/ERT2)MRhn/CdnyJ (ArcCreER^T2^) mice were crossed with B6.129X1-Gt(ROSA)26Sor^tm1(EYFP)Cos^/J (eYFP) mice (Jackson laboratory strain #022357 and strain #006148, respectively) to produce double transgenic mice (hereafter referred to ArcCreER^T2^ x eYFP mice). ArcCreER^T2^ x eYFP mice were bred on-site over multiple generations. At the memory retrieval stage of all experiments, adults ranged from 71 – 150 days old (mean = 112.5 ± S.D. 27.8) (female weight = 16.4 – 32.9 g, median weight = 22.1; male weight = 21.9 – 42.8 g, median weight = 30.4). Same sex mice were group-housed (2-5/cage) in individual vented standard cages, except when singly housed after surgery. There were three types of enrichment in each cage, including a wood gnawing block, nestlet, and EnviroPAK. The vivarium temperature (23 - 25 °C), humidity (35 - 37%), and 12 hr light/dark cycle (lights on 0600) were controlled throughout. Food and water were available *ad libitum* and cages were changed 1 x/week. Mice were randomly assigned to experimental groups based on litter before the start of each experiment. All experimental procedures were conducted in accordance with the National Institutes of Health guidelines on the Care and Use of Animals in Research and approved by the Vassar College Institutional Animal Care and Use Committee (IACUC). Disclosure of animal housing, husbandry, and experimental procedures follow principles for transparent reporting and reproducibility in behavioral neuroscience (Prager et al., 2011, 2018).

### Genotyping protocol

Tissue biopsy was performed by tail snip under brief isoflurane anesthesia. DNA was extracted using DirectPCR Lysis Reagent (Viagen Biotech, Los Angelas CA) following manufacturer protocol. Amplification of Cre and R26R was performed using the following primer sets: *Cre*: 5′-GCC TGC ATT ACC GGT CGA TGC AAC G-3′; 5′-AAA TCC ATC GCT CGA CCA GTT TAG TTA CCC-3′. *R26R* (for *EYFP* mice) 5′-GGA GCG GGA GAA ATG GAT ATG-3′; 5′-AAA GTC GCT CTG AGT TGT TAT-3′; 5′-AAG ACC GCG AAG AGT TTG TC-3′, following recommended cycling protocols from Jackson Labs and Denny et al., 2014.

### General behavioral experimental procedures

All experiments were performed during the light cycle. A background strain of the ArcCreER^T2^ x eYFP mouse is the C57BL/6J (B6) mouse, which is an age-related hearing decline model (Ison et al., 2007). Hearing decline in B6 mice in complex, but low frequency hearing, which is relevant to this study, has consistently been shown to decline with age. To address this, no mice over 150 days old were tested. We also evaluated peripheral auditory brainstem responses to relevant frequencies (see below). All mice were habituated to a holding room 30-45 min prior to conditioning and testing. To reduce contextual (background) freezing, the training context (hereafter referred to as “Context A”) was disguised from the testing context (hereafter referred to as “Context B”) using several manipulations. Context A was an unmodified fear conditioning chamber (Coulbourn Instruments, Holliston, MA) and a 70% EtOH solution was used to clean the chambers between mice. For context B, (1) mice were transferred from the vivarium to the holding room using distinctive cages, carts, and covering, (2) the ambient lighting and background noise of the holding and testing room were changed using different illuminance and a fan for background noise, (3) a white plexiglass floorboard sprinkled with clean bedding was used to cover the shock bars, (4) the testing chamber walls were disguised with black- and white-striping, (5) the chambers were cleaned with a 1% acetic acid solution between mice (Bergstrom, 2020). Each day, prior to training and testing, the decibel level (dB) for the auditory tone frequency was measured in each chamber using a sound level meter (R8050, REED Instruments, Wilmington, NC) and calibrated to 70-75 dB. All conditioning was conducted in commercial chambers (20 x 30 x 18 cm) in sound-dampening cabinets (58 x 61 x 45 cm) (Colbourn instruments, Holliston, MA). FreezeFrame 4 software was used for controlling and delivering the tone and foot shock stimuli (ActiMetrics, Wilmette, IL). All acoustic stimuli were delivered by a speaker mounted on the upper center of one wall and foot shock stimuli were delivered via a stainless-steel rod floor (0.8 cm distance between rods, 1.0 cm rod diameter).

### Fear conditioning

Mice were placed in the fear conditioning chamber (Context A) 180 s prior to 3 pairings of an auditory tone CS (20 s, 5-kHz, 70-75 dB) that co-terminated with an electric foot shock US (0.5 s, 0.5 mA). The CS/US pairings were separated by variable inter-trial intervals (ITI) (20 and 80 s). Mice were removed from the chamber 60 s after the final CS/US pairing. The total conditioning time was 400 s.

### Context B pre-exposure

To reduce “background” generalized contextual freezing (Jacobs et al., 2010), mice were placed into Context B for 15 min and left to explore on the day prior to the cued generalization test (Bergstrom, 2020).

### Generalization test

Either 6 days (recent group) or 30 days (remote group) following training (1 day following Context B pre-exposure) mice were placed in Context B prior to 3 presentations either the CS (5-kHz, 70-75 dB, 20 s) or a novel tone generalization stimulus (GS; 3-kHz, 70-75 dB, 20 s). A 3-kHz tone GS was used because in previous experiments, following conditioning with a 5-kHz CS, it produced the greatest degree of generalization over time in comparison with alternate tone frequencies (Pollack et al., 2018). The novel 3-kHz tone GS was dubbed the “ambiguous” stimulus throughout. Mice were removed from the chamber 60 s after the final stimulus presentation and returned to the colony room (400 s total test time; ITIs 80 and 20 s). Mice in the “no tone” control group were allowed to explore context B for 400 s.

### Genetic labeling procedures

To reduce non-specific genetic labeling, mice were dark-housed the night before and three days following the 4-OHT injection. These methods have been previously validated (Cazzulino et al., 2016; Denny et al., 2014). For dark housing procedures, mice were placed in a separate housing room with lights off, limited noise, and no handling. ArcCreER^T2^ X eYFP mice were intraperitoneally (i.p.) injected with 4-OHT exactly 5 h prior to fear conditioning. This timeframe has been previously validated (McGowan et al., 2024).

### Drugs

#### 4-Hydroxytamoxifen (4-OHT)

Cre-mediated recombination in ArcCreER^T2^ x eYFP mice was induced using 4-OHT (HelloBio, Princeton NJ; SKU: HB6040). 4-OHT was made fresh prior to each injection (i.p.). 4-OHT was dissolved by water bath sonication in a 10% EtOH/90% corn oil solution at 10 mg/mL. The final dose was 55 mg/kg.

### DREADD actuators

#### Clozapine N-oxide (CNO)

For the CaMKII-DREADD experiments, CNO was used as the chemical actuator (Hello Bio, Princeton NJ SKU: HB6149). It was made fresh prior to i.p. injection, dissolved in 0.9% saline (1 mg/mL), and injected i.p. (5.0 mg/kg) exactly 45 min prior to the memory test.

#### Deschloroclozapine dihydrochloride (DCZ)

For DIO-DREADD experiments, DCZ was used as the chemical actuator (Nagai et al., 2020) (Hello Bio, Princeton, NJ, SKU: HB9126). It was made fresh immediately prior to i.p. injection, dissolved in 0.9% saline (1 mg/mL), and injected i.p. at 100 µg/kg exactly 15 min prior to the memory test.

### Immunohistochemistry

#### Tissue collection

Exactly 90 min following the fear memory retrieval test, all mice were injected with a ketamine/xylazine cocktail (100:10 mg/mL) and transcardially perfused with ice-cold 1X PBS (7.2 - 7.4 pH), followed by ice-cold 4% paraformaldehyde (PFA) in 1X PBS (7.2 - 7.4 pH). The time point for perfusion following the retrieval test was based on several previous reports (Maddox and Schafe, 2011; Ploski et al., 2008). Brains were extracted and placed into 4% PFA overnight at 4°C then transferred to 1X PBS and kept at 4°C until sectioning. Coronal brains sections (40 μm thick) were cut through the mPFC using a vibratome (VT1200, Leica Biosystems Inc., Buffalo Grove, IL). Every other section (to avoid double-counting) was collected in a well plate containing 1X PBS (7.4 pH) for free-floating immunohistochemistry. Sections were rinsed first in 1X PBS (3 x 10 m), blocked in a 1X PBS/1% bovine serum albumin (BSA)/0.2%Triton-X solution for 1 hour, and incubated in antigen specific antibodies (see below). All incubations were performed on orbital shakers.

#### Arc and eYFP primaries

After blocking, sections were incubated overnight in Anti-Arc rabbit polyclonal antibody (1:5000) (Cat No. 156003, Synaptic Systems, Goettingen, Germany) and Anti-GFP chicken polyclonal antibody (1:10000) (Cat No.13970, Abcam, Waltham, MA) at 4°C. The next day, sections were washed in 1X PBS (3 x 10 m) before a 1 h incubation in Donkey anti-rabbit IgG (1:1000) (#A32754, Invitrogen) and Goat anti-chicken IgG (1:500) (Cat No. A32931, Invitrogen) at room temperature.

#### c-Fos

To test the efficacy of CNO to activate the DREADDs system (either hM4Di or hM3Dq), a subset of mice, having completed all behavioral testing, received i.p. injections of CNO at 5.0 mg/kg dosage and were placed into the Context B recall test after 45 min (peak expression time) (Campbell & Marchant, 2018). Exactly 135 min after CNO injection and 90 min after recall (peak c-Fos expression) (Barros et al., 2015), mice were sacrificed for IHC. All procedures for IHC were identical to those described above except that sections were then incubated for 24 hours with anti-c-Fos rabbit polyclonal antibody (1:500) (Cat No. RPCA-c-Fos, RRID: AB_2572236, EnCor Biotechnology Inc., Gainesville, FL USA) at room temperature. For the secondary antibody, subjects in either the pAAV-CaMKIIa-hM4Di-mCherry or pAAV-CaMKIIa-hM3Dq-mCherry groups were incubated with Alexa Fluor 488 goat anti-rabbit IgG (H+L) (Lot #: 1981125, REF: A11008, Invitrogen, Carlsbad, CA USA) and pAAV-CaMKIIa-EGFP subjects were incubated with Alexa Fluor 594 donkey anti-rabbit IgG (H+L) (Lot #: WD319534, REF: A32754, Invitrogen, Carlsbad, CA USA) for one hour at room temperature.

### DIO-DREADD Arc

To test the efficacy of DCZ to activate the DIO-DREADDs system (either DIO-hM4Di or DIO-hM3Dq) and modify Arc expression in preferentially transfected cell populations, mice were injected with DCZ 105 min prior to fear conditioning and processing for Arc IHC. The timeline for DCZ injection were based on high DCZ brain level concentrations at 15 min (Nagai et al., 2020) and peak Arc expression at 90 min following fear conditioning (Maddox and Schafe, 2011; Ploski et al., 2008). The experimental procedures for IHC were identical to those describe above for Arc IHC except the secondary antibody for Arc was Donkey anti-rabbit IgG (Alexa Fluor 647) (1:1000) (Lot #: YF374181, REF: A32795, Invitrogen, Carlsbad, CA USA).

#### Tissue mounting

Following all incubations, sections were rinsed a final time in 1X PBS (3×10 m) then mounted onto gel-coated slides in 0.05M PB. Sections were cover slipped (#1 or #1.5 thickness) using Fluoromount-G mounting medium with DAPI (#00-4959-52, Invitrogen) and sealed with nail polish. Slides were stored at 4°C in darkness until imaging.

#### Image acquisition and analysis

Images were acquired using a Leica TCS SP5 II laser scanning confocal with the Leica Microsystems LAS AF software (Version: 2.6.0.7266). The objective used was a Leica HCX PL APO CS 20X/.70 Dry. 3D images were acquired by taking a z stack of 20-30 slices with 1.14 µm spacing and pixel dimensions 760 x 760 nm. Images for the DIO-DREADD experiment were captured using the Leica Stellaris 8 FALCON laser scanning confocal platform.

The experimenter was blind to experimental conditions throughout all neuronal ensemble quantification procedures. Cell counts were acquired by sampling across 6 IL regions/subject. Both hemispheres were included in the analysis. IL sampling regions for data collection were chosen based on the quality of the staining and visibility of anatomical landmarks for localization of the counting frame (see below). Labeled cells were quantified using FIJI. Background subtraction was applied across all channels. For c-Fos microscopy, images were captured under fluorescent microscope (Nikon Eclipse 50*i*, Nikon Instruments, Amsterdam, NL). c-Fos^+^ cells were counted manually (FIJI-ImageJ open source) in the mPFC using a counting frame (250 x 250 *µ*m) with AAV^+^ and adjacent AAV^-^ expression area and six locations/subject (mCherry^+^ n = 6, eYFP^+^ n = 6).

To locate IL and PL for placement of the counting frame, the lateral ventricle (LV) was used as an anatomical landmark, as it is readily identifiable and located in a consistent position ventral to the IL, and aligned with DP, for a majority of the longitudinal axis. Midline and the corpus collosum were also used as stable anatomical landmarks to identify the location of the IL and PL. For the IL, the counting frame (250 x 250 *µ*m) was positioned approximately 600 *µ*m dorsal from the LV. For the PL, the counting frame was positioned 1250 *µ*m dorsal from LV. For layer measurements, the counting frame was centered 300 *µ*m from midline for L2/3 (shallow layers) and 500 *µ*m (deep) from midline for L5/6. Cells in the IL were counted between rostrocaudal levels of bregma 1.93 and 1.53 (Paxinos & Franklin, 2019). DAPI cells were counted using the 3D object counter in FIJI. Fluor 488+, Fluor 594+, and co-labeled cells were counted manually within the counting frame. Colocalized eYFP^+^:Arc^+^ neurons were first analyzed as a percentage of the number of DAPI^+^ neurons. Chance rate neuronal co-localization was calculated as: (eYFP^+^/DAPI^+^) * (Arc^+^/DAPI^+^) * 100.

### DREADDS

#### Viral Constructs

All viral constructs were purchased from Addgene (Watertown MA). In the first set of experiments, pAAV-CaMKIIa-hM4Di-mCherry (AAV5), pAAV-CaMKIIa-hM3Dq-mCherry (AAV8), or a fluorophore-only control AAV (pAAV-CaMKIIa-EGFP; AAV5) was used. In the DIO-DREADD experiments, pAAV-hSyn-DIO-hM4D(Gi)-mCherry (AAV8), pAAV-hSyn-DIO-hM3D(Gq)-mCherry (AAV8), or fluorophore control AAV (pAAV-hSyn-DIO-mCherry (AAV8) was used. All viral vectors were aliquoted and stored at −80 C until use.

#### Surgery

Prior to surgery, mice received an injection of carprofen (s.c., 5.0 mg/kg) and an intradermal injection of bupivacaine (0.05 mL) at the craniotomy site. Inhaled isoflurane levels were maintained between 1.25% - 2.5% throughout. Mice were bilaterally microinjected (100-150 nL/hemisphere) with either the hM4Di, hM3Dq, or fluorophore-control AAV targeting the IL (stereotaxic coordinates: AP: +1.8, ML: +0.3/-0.3, DV: −2.8). Microinjections were conducted using a 2.5 µL glass syringe and 32-gauge needle (Hamilton, Reno NV). Following surgery, a dietary supplement (DietGel Recovery Purified) and carprofen (s.c., 5.0 mg/kg) was provided as needed. Subjects were singly housed for two weeks and group-housed, if possible, prior to behavioral testing. Fear conditioning occurred no less than two weeks following stereotaxic surgery. All fear conditioning and generalization experimental parameters were identical to those described above.

On generalization test day, mice were injected (i.p.) with CNO 45 min prior to testing. The following day, mice were injected (i.p.) with saline (same volume as CNO) using procedures identical to those described above for CNO and were run through the generalization test protocol again.

### DIO-DREADD methods

To manipulate neuronal ensemble reactivation, a Cre-dependent viral construct expressing either an excitatory DREADD (pAAV-hSyn-DIO-hM3D(Gq)-mCherry), inhibitory DREADD (pAAV-hSyn-DIO-hM4D(Gi)-mCherry) or a fluorophore control (pAAV-hSyn-DIO-mCherry) was injected into the IL of ArcCreER^T2^ x eYFP mice. After at least 2 weeks, on the fear conditioning day, all mice were injected with 4-OHT to drive Cre-recombination and DIO-DREADD expression in neurons with high levels of Arc. This permits specific expression of DREADDs constructs in activated cellular populations for later reactivation or silencing using the actuator DCZ.

Either 6 days (recent group) or 30 days (remote group) following training (1 day following Context B pre-exposure), mice were injected with DCZ 15 min prior to the generalization test. 90 min following testing, all mice were sacrifice for Arc IHC (see above).

### Behavioral analysis

#### Freezing

An overhead camera recorded digital video of the fear conditioning chamber. Freezing behavior was automatically quantified using FreezeFrame 4.0 (ActiMetrics, Wilmette IL). Freezing was defined as the lack of movement except for respiration for >1 s (FreezeFrame threshold = 5). Percentage freezing data was calculated by scoring freezing during the CS or GS presentations (20 s), pre-CS/GS period (habituation), and inter-trial intervals (ITIs).

#### Scanning

Scanning was operationally defined as a side-to-side head and front paw movement while the tail base remained motionless. Scanning behavior was recorded immediately upon initiation and halted when the mouse, 1) froze >1 s or, 2) initiated full movement. Movement was defined as a larger category of behaviors, including four-paw locomotion, grooming, rearing, and small, jerk-like body and head movements, not identified as freezing or scanning.

Pose estimation for each video was created using DeepLabCut version 2.3.10. Each video was 400 s long and recorded at 8 frames/s (fps), resulting in 3,000 frames/video. Fifty randomly sampled frames from each video were manually labeled, with the nose and tail base serving as the labeling points.

To train the ResNet-50 network, 95% of the labeled frames were used, while the remaining 5% were reserved for testing the neural network’s performance. Each video underwent over 10,000 training iterations, yielding a training error of 8.02 pixels and a test error of 8.61 pixels. By applying a p-cutoff of 0.4, the training error was reduced to 7.22 pixels, though the test error remained unchanged.

A separate Python script was developed to extract freezing and scanning behaviors from DeepLabCut (DLC) output, which included pose estimations of nose and tail base. This script averaged the coordinates over 8 frames to determine the coordinates/s. It then calculated the velocity (pixels/s) of nose and tail base/s. The script identified freezing and scanning behaviors using criteria established through extensive comparisons between test results, hand-scored, and Freezeframe 4.0 results (ActiMetrics, Wilmette IL). In DLC, freezing behavior was defined as the velocity of both the nose and tail base being < 6 pixels/s with immobility lasting >1 s. Scanning behavior was defined as the nose velocity >10 pixels/s, and tail base velocity <10 pixels/s.

### Auditory Brain Stem Responses

To determine auditory thresholds, a set of 5-ms tonebursts (1-ms Blackman-Harris gating) was generated in SigGenRZ (v 5.6.0). We generated stimuli at nine frequencies (1, 2, 2.5, 3, 3.15, 4, 8, 10, and 12.5 kHz) that spanned the frequency range of stimuli used in the generalization experiments. We also generated broadband clicks that were periodically presented to the animals to track physiological stability across the course of the experiment. Peak-to-peak equivalent stimuli levels were determined with a Larson Davis LXT sound level meter and a long-duration 1 kHz tone. We calibrated the frequency response of the speaker with long-duration tones in 1/3 octave bands with a Larson Davis LXT sound level meter (fast, z-weighting). Frequency-specific output levels were adjusted using the gain function in SigGenRZ until a flat frequency response was achieved (± 1 dB).

All tests were performed in a 1.8 m x 1.9 m x 2 m IAC acoustics (Naperville, IL) audiology booth lined with pyramidal acoustic foam to provide sound deadening. We used electrode placement and stimulus presentation rates previously used. Each subject was anesthetized with an injection (i.p.) of ketamine (90 mg/kg) and xylazine (10 mg/kg) solution. The subject was placed on a heating pad covered with surgical towels. When the animal no longer responded to a toe-pinch, we cleaned the skin with 70% isopropyl alcohol and three 27-gauge 12 mm subdermal needles (Rochester Electro-Medical Inc.; Lutz, FL) were inserted: one non-inverting (active) electrode at the vertex of the head, one inverting (reference) electrode directly below the auditory meatus of the right ear, and one grounding electrode directly below the auditory meatus of the left ear. The electrode leads were connected to a Tucker Davis Technologies (TDT; Alachua, FL) RA4LI head stage and RA4PA preamp, which then fed into a TDT RZ6 processor via a fiber optic cable. After placing the electrodes, the impedance of the electrode was checked and repositioned if necessary to maintain an impedance at or below 5 kΩ. During the experiment layers of surgical towel were periodically added or removed to maintain body temperature.

Stimulus presentation and evoked potential recordings were coordinated by BioSigRZ (v 5.6.0), a POE5 signal processing card, and the RZ6 processor. Stimuli were presented at a rate of 31.1 stimuli s-1 from an Orb Mod2 satellite speaker (Orb Audio, U.S.A.; frequency response: 0.12 – 15 kHz) positioned 10 cm from the right ear of the subject. Different stimulus amplitude intervals were used depending on known thresholds for each frequency to assess thresholds more rapidly. Typically, larger steps (10 or 20 dB) were used farther away from the threshold and smaller steps (5 dB) were used near the threshold. The exact set of stimuli amplitudes varied by frequency. Two sets of 400 stimuli were played in alternating phases for each combination of stimulus frequency and amplitude. Between each set of frequencies, we also assessed the response to a pair of clicks presented at 80 dB to monitor the physiological stability of the subject. Evoked responses were notch-filtered at 60 Hz and band-pass filtered between 0.03 – 3 kHz.

Auditory thresholds at each frequency were determined using visual detection, where two trained observers independently identified the lowest stimulus amplitude evoking a response. Thresholds were estimated as the sound pressure level halfway between that of the last detectable response and the next quietest stimulus. Since stimulus intensities in threshold regions differed by 5 dB, ABR thresholds were defined as the intensity 2.5 dB below the lowest stimuli amplitude at which a response could be visually detected.

### Experimental Design and Statistical Analysis

For all behavioral data, a mixed ANOVA was used to compare the conditioned freezing response across CS/GS and ITI presentations (within-subject variable) across groups. Male and female mice were included in all experiments in a full factorial experimental design. All data were first checked for normality using Mauchley’s test for sphericity. Violation of the assumption of sphericity was addressed by adjusting the degrees of freedom using the Greenhouse-Geisser correction. For some analyses, a generalization index was calculated and compared across groups. The generalization index was calculated by dividing the mean of the CS by the sum of CS and GS [CS / (CS + GS)]. Although the generalization index range is 1 to 0, a value of 1 indicates no generalization and a value of 0.5 indicates complete generalization.

For the ArcCreER^T2^ x eYFP tagging experiments, a multivariate analysis of variance (MANOVA) was used to test the statistical relationship among eYFP^+^, Arc^+^, or eYFP^+^:Arc^+^ co-labeled cells in L2/3 and L5/6 of the PL and IL at different kHz frequencies (0-kHz control, 3-kHz, 5-kHz) and time points (Recent or Remote) following learning. Statistics were run on the number of eYFP^+^ / DAPI^+^, Arc^+^ / DAPI^+^, eYFP^+^:Arc^+^ / DAPI^+^ neurons compared with the chance rate of co-activation (eYFP^+^ / DAPI^+^) * (Arc^+^ / DAPI^+^) * 100), and the number of eYFP^+^ : Arc^+^ / DAPI^+^ neurons minus chance rate (Reijmers et al., 2007; Tayler et al., 2013). Box’s M test was used to test the equality of variance-covariance matrices. Violation of the homogeneity assumption was followed up by a rank order transformation. A significant value for the conservative Pillai’s Trace test statistic was only followed up by Bonferroni-corrected univariate ANOVAs. Follow-up ANOVAs were checked for the assumption of equality of covariance matrices using Levine’s test. The Welsch test was used in the case of a significant Levine’s test. A significant ANOVA was followed up with a Scheffe post hoc test. Prior to analysis, outliers were determined by calculating the interquartile range for each group. Any values > 1.5 steps beyond the interquartile range were considered outliers and removed from the analysis. These are reported in the results. For all statistics, significance was set at *p*<.05. All data are represented as the mean ± the standard error of the mean (SEM). All group sizes can also be found in the figure captions. Group sizes were based on previous studies (Pollack et al., 2018; Scarlata et al., 2019). Statistics were run on SPSS (IBM, Armonk, NY v. 26).

## RESULTS

### Experiment 1: Bidirectional IL chemogenetic control during the expression of cue fear memory generalization

#### DREADDs system efficacy

Microinjections of the stimulatory DREADD (hM3Dq), inhibitory DREADD (hM4Di), or fluorophore control were administered into IL at least 2 weeks prior to behavioral testing or c-Fos analysis (Figure 1A). To assess the efficacy of the DREADDs system, we analyzed mPFC c-Fos^+^ cell density after both hM3Dq and hM4Di CNO-induced stimulation. Results showed robust CNO-induced effects across DREADDs constructs (*F*[2, 13]=38.8; *p*<.001), with increased (*p*<.001), and decreased (*p*<.01), c-Fos^+^ cell density relative to the fluorophore control, respectively (Figure 1B and C). All injections were stereotaxically biased towards vmPFC coordinates to avoid transfection in the dorsal medial prefrontal cortex (dmPFC). After exclusion of mice with dmPFC transfection, histological analysis mapping the extent of viral transfection across individually aligned brains in stereotaxic group space confirmed AAV transfection predominantly located to IL, with some expression in dorsal peduncular cortex (PD) (Figure 1D).

**Figure 1.**
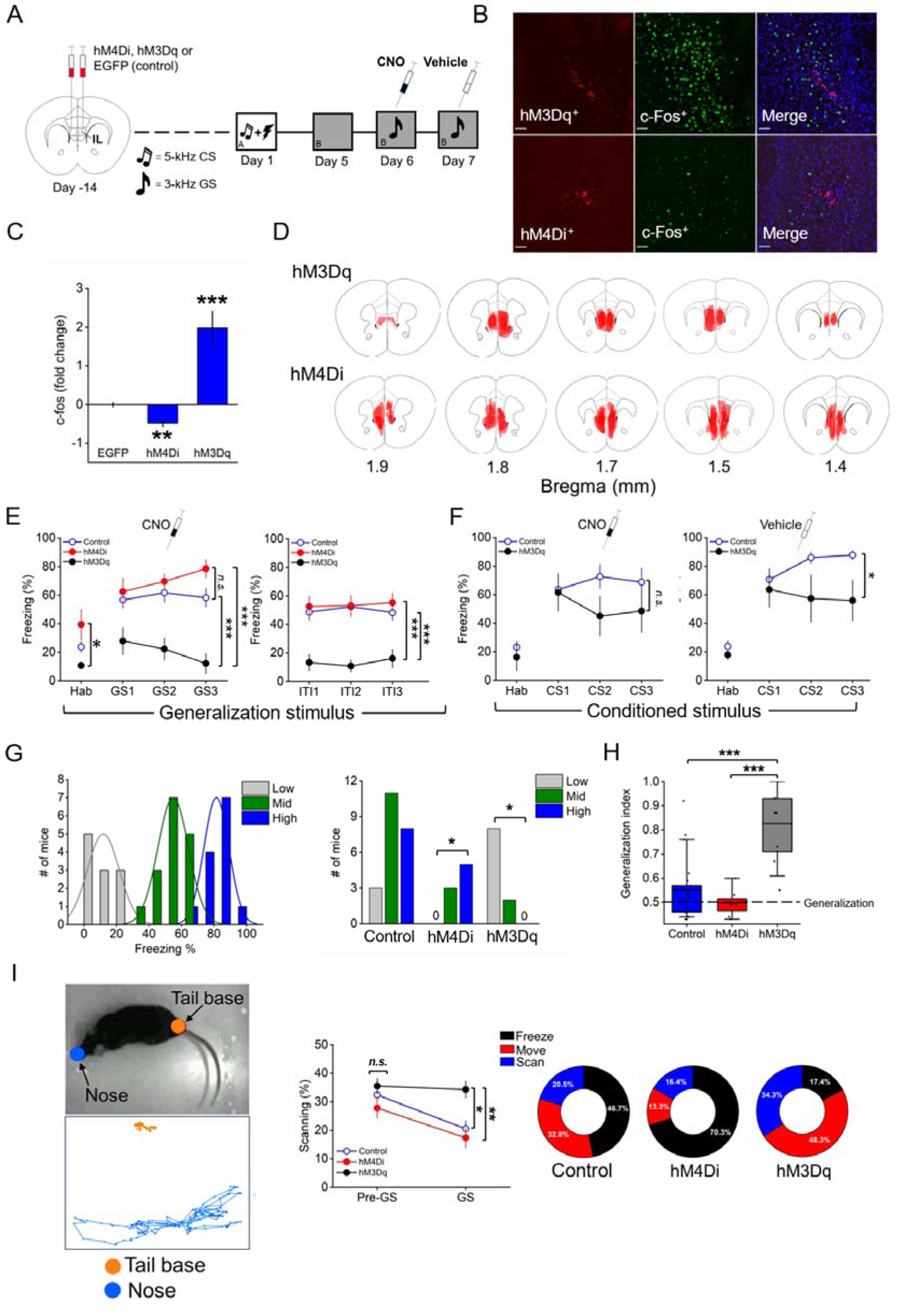
Bidirectional chemogenetic IL manipulation during fear generalization expression. (A) Experimental design. (B) Representative confocal micrographs showing AAV^+^ expression (left panel), c-Fos^+^ FIH (middle panel), and AAV^+^/c-Fos^+^/DAPI^+^ co-expression (20X/0.8 N.A. air). Scale bar = 25 *µ*m. (C) There was a 2-fold increase in c-Fos^+^ expression in hM3Dq mice (n=4) and nearly 0.5-fold decrease in c-Fos^+^ expression in hM4Di mice (n=6) versus controls (n=6). (D) The extent of AAV expression in the mPFC across five coronal planes is depicted. Relatively darker shaded regions show greater overlap across individuals. Transfection was predominantly located in the IL, with some expression in DP. Mouse brain atlas images modified from (Paxinos & Franklin, 2004). (E) IL stimulation (hM3Dq; n=10) reduced freezing to the context, GS stimulus, and during the intertrial interval (ITI) compared with hM4Di mice (n=8) and fluorophore controls (n=22). (F) In a separate group of mice, there was no CNO-induced differences between hM3Dq (n=5) and control (n=12) mice in response to CS presentation. The next day, after vehicle injection there was reduced freezing in the hM3Dq group. (G) An unbiased k-means clustering algorithm run on all mice produced 3 prominent clusters (Low, Mid, and High). Chi-square analysis indicated all mice in the DIO-hM4Di were clustered in the “High” and “Mid” group and a majority of mice in the DIO-hM3Dq groups clustered in the “Low” group. (H) Analysis of the generalization index showed greater values (less generalization) in the hM3q group versus the hM4Di group and controls. (I) (Left) DeepLabCut analysis of scanning behavior. (Top panel) markers for pose estimation on the tail base and nose and (bottom panel) representative path analysis depicting low activity in tail base and high activity in nose (scanning). (Middle) Prior to GS presentation, there was no differences in scanning behavior between groups. Upon presentation of the GS, scanning collapsed in the DIO-hM4Di group but was maintained in the DIO-hM3Dq group. (Right) Donut plots depict percentage freezing, scanning, and moving during the mean GS presentations. n=8-22/group \**p*<.05 \*\**p*<.01, \*\*\**p*<.001, n.s., non-significant.

#### Fear generalization test during IL manipulation

CNO was injected prior to a cued fear memory generalization test to activate the DREADD construct. Results revealed a main effect of the DREADD manipulation (*F*[2, 37=18.9; *p*<.001). In the hM3Dq group, both GS and ITI-elicited freezing (ITI data not shown) was reduced compared with the control (*p*<.001) and hM4Di group (*p*<.001) (Figure 1E). There was also a smaller, but significant, effect of the DREADD manipulation on pre-CS freezing (*F*[2, 37=4.1; *p*=.024). hM3Dq mice froze less than hM4Di (*p*=.027). These data indicate IL excitatory stimulation suppresses the expression of auditory, and to a lesser extent, contextually conditioned generalized freezing. On the next day, without CNO (vehicle injection), there was no difference in tone-elicited freezing between groups (data not shown), further supporting data showing IL excitatory manipulation impacts generalized freezing responses. There were no sex differences detected throughout.

#### IL stimulation, conditioned freezing, and extinction

Because IL stimulation consistently decreased freezing in response to the GS, an important consideration is the generality of IL function in suppressing conditioned freezing responses. To test this question, we replicated the generalization protocol described above in a separate set of hM3Dq and fluorophore control mice, but rather than presenting the GS, we presented the CS. Results showed no difference in CS-eliciting freezing between hM3Dq and controls group at the first CS and throughout CS presentations (Figure 1F). On the following day, the same hM3Dq mice injected with vehicle showed a modest, but significant, decrease in CS-elicited freezing as compared with the fluorophore control (*p*=.042), suggesting facilitation of extinction consolidation.

In our paradigm, and especially in these initial experiments, presentation of GS produced relatively high levels of generalized freezing (60.5%) and relatively high variance (SD: 23.5). This finding led us to examine individual differences in generalized responses. An unbiased k-means clustering algorithm was applied to mean GS freezing levels across all mice. The analysis revealed 3 prominent “High, Mid, and Low” clusters (Figure 1G). A reanalysis of the fear conditioning acquisition data showed no differences between low, mid and high groups, indicating this behavioral phenotype is not present during learning. A chi-square test was performed to test the relationship between experimental groups and clusters. Results showed a significant effect ^2^ (4, N=40) = 20.9, *p*<.001, with mice in the hM4Di group preferentially clustered in a rightward skewed direction in “High (n=4)” and “Mid (n=3)” generalization groups with none in the “Low” group. Mice in the hM3Dq group clustered leftward almost exclusively in the “Low (n=8)” generalization group (2 mice clustered in the “Mid” group) with none in the “High.” This is in contrast to the control mice that displayed a more normally shaped distribution across the “Low,” “Mid,” and “High” groups. This unbiased analysis further confirms IL silencing drives more generalized freezing responses, while IL stimulation promotes less generalized freezing. In a final analysis of the generalization index, results showed a significant effect of the DREADDs manipulation (F[2, 34] = 4.7; *p*<.001) (Figure 1H). Mice in the hM3Dq groups exhibited greater scores (less generalization) relative to the hM4Di (*p*<.001) and Control group (*p*<.001). As a further control, we analyzed data obtained from mice with unilateral DREADDs expression in IL. Surprisingly, we found no effect of unilateral hM3Dq or DREADDs activation on generalization expression.

#### IL stimulation and scanning behavior

Next, we asked the question, if hM3Dq mice exhibited reduced freezing to the GS how were they behaving? We speculated that IL stimulation might promote movement classified under “threat detection” (Blanchard et al., 2011), or “pre-encounter threat responses” (Roelofs & Dayan, 2022), such as scanning behavior (Choy et al., 2012). To test this hypothesis, scanning behavior was analyzed using machine learning technology (DeepLabCut2.0) (Mathis et al., 2018; Nath et al., 2019) and custom code. RMANOVA showed a Time x Group interaction (F[2, 35]=3.7; *p*=.036). There were no differences in scanning behavior prior to the presentation of the GS across all groups (Figure 1I). However, upon GS presentation, scanning behavior collapsed in the hM4Di (*p*=.01) and control group (*p*=.014), but was maintained in the hM3Dq (*F*[2, 35]=6.5; *p*=.004) (Figure 1I). These data indicate a role for IL in suppressing freezing and maintaining scanning. Interestingly, scanning behavior made-up approximately 50% of all movement across all groups, regardless of DREADDs manipulation. Overall, IL excitatory activity is sufficient to suppress post-encounter defensive responses (freezing) in favor of pre-encounter defensive (scanning) and non-defensive (movement) behaviors in response to an “ambiguous” threat stimulus.

#### Brainstem auditory evoked potentials

A fundamental question in the study of stimulus generalization is the consideration that generalized responses may represent a failure in perceptual (sensory) discrimination (Dunzmoor and Paz, 2015), rather than mnemonic processes per se (Zaman et al., 2021). This might confound the present results indicating generalization at the level of forebrain plasticity. To begin to address this question, auditory thresholds using auditory brainstem responses were tested in a subset of ArcCreER^T2^ x eYFP mice. While thresholds cannot determine whether two tones are discriminable from one another, they can address whether tones are likely to be detectable. Evoked potential threshold estimates tend to underestimate behavioral thresholds, so stimuli presented above the AEP threshold should be of sufficient amplitude to evoke behavioral responses. The threshold by frequency response we found is consistent with the patterns in C57BL/6J (B6) mice. Generally, thresholds decreased (i.e. sensitivity improved) as stimulus frequency increased, although the rate of change increased with each subsequent octave. Thresholds at 3 kHz ranged from 57.5 to 67.5 dB SPL, with an average of 62.5 dB SPL. These thresholds are lower than the amplitude of the stimulus used in the generalization experiments, suggesting the ArcCreER^T2^ x eYFP mice should be able to detect this tone. Thresholds do decrease between 4- and 8-kHz, suggesting that the 5-kHz tone may have a greater sensation level than a tone at 3-kHz.

Estimates of auditory filter bandwidth using auditory evoked potentials or psychophysical frequency discrimination limens would further address whether individuals from this strain of mice differ in their ability to discriminate between these tones. C57BL/6J (B6) mice are capable of discriminating frequency changes of 250-Hz or less at 8-kHz, with frequency discrimination decreasing with increasing frequency. Absolute frequency discrimination limens typically decrease with decreasing frequency (Weber’s law), suggesting mice should have no difficulty discriminating at 3-kHz tone from a 5-kHz tone. Although sensitivity is lower at 3-kHz than at 5-kHz, our results suggest that tones of both frequencies should be detectable at 70-75 dB in our subjects.

### Experiment 2: Cued fear memory generalization and the passage of time

Here we tested whether cued fear memory generalization increases over time in ArcCreER^T2^ x eYFP transgenic mice, a finding previously shown in the C57BL/6N substrain (Pollack et al., 2018). All mice were fear conditioned with the 5-kHz CS and then, either 7 days (recent) or 30 days (remote) later, were presented with either the CS again or the GS. At the recent time point, there was less freezing in response to the GS (3-kHz) compared to the CS (5-kHz) (*F*[1,40]=13.4; *p*<.001) (Figure 2B).

**Figure 2.**
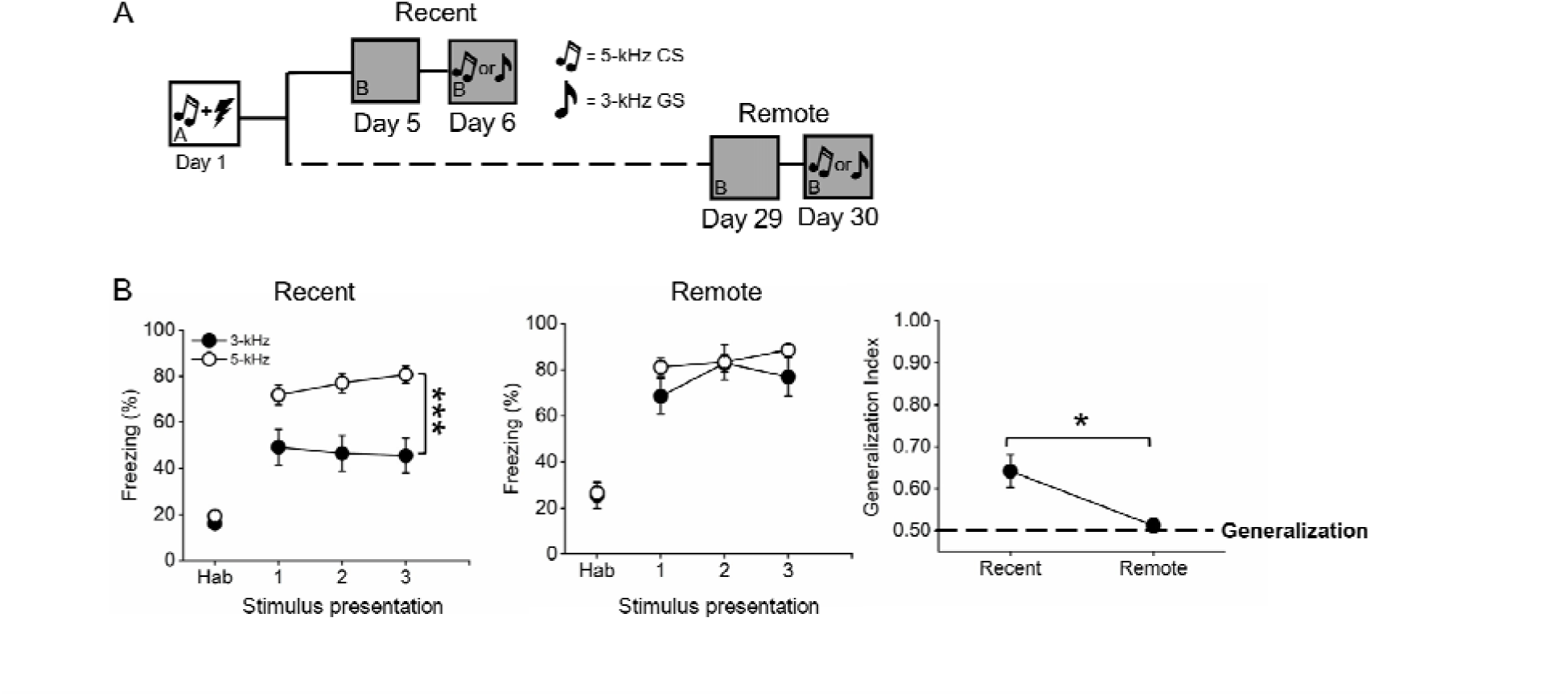
Cued fear memory generalization over time. (A) Schematic of the experimental design. (B) At the recent timepoint following learning (7 days), mice in the 3-kHz group exhibited less freezing than the 5-kHz group. At the remote timepoint (30 days), freezing was equivalent between 3-khz and 5-kHz groups. The generalization index reduced over time, indicating greater generalization at the remote compared with the recent timepoint. n=9-21/group. \**p*<0.05, \*\*\**p*<0.001.

At the remote time point, there were no differences between the GS and CS groups (*p*=.58). Analysis of the generalization index showed a modest, but significant, reduction over time (*F*[1, 28]=4.3; *p*=.045; Figure 2B), indicating enhanced generalization with the passage of time, a finding congruent with previous work (Pollack et al., 2018).

### Experiment 3: mPFC neuronal ensemble quantitative measures

In this experiment, a new cohort of mice underwent experiments described in Experiment 2, but 90 min after generalization tests, brain tissue was harvested and processed for Arc immunohistochemistry (IHC). A “no tone” control group was included as a baseline measure in which all procedures were identical to the experimental groups, except that on the test day, mice were not presented with a tone but instead left to explore the context for the same amount of time as the tone stimulus groups (Figure 3A). mPFC neuronal ensembles were tagged using the ArcCreERT2 x eYFP system in which 4-OHT administration induces Cre-mediated recombination and expression of the eYFP reporter in activated neurons (Figure 3B-E). Neurons were counted in PL and IL (Figure 3F).

**Figure 3.**
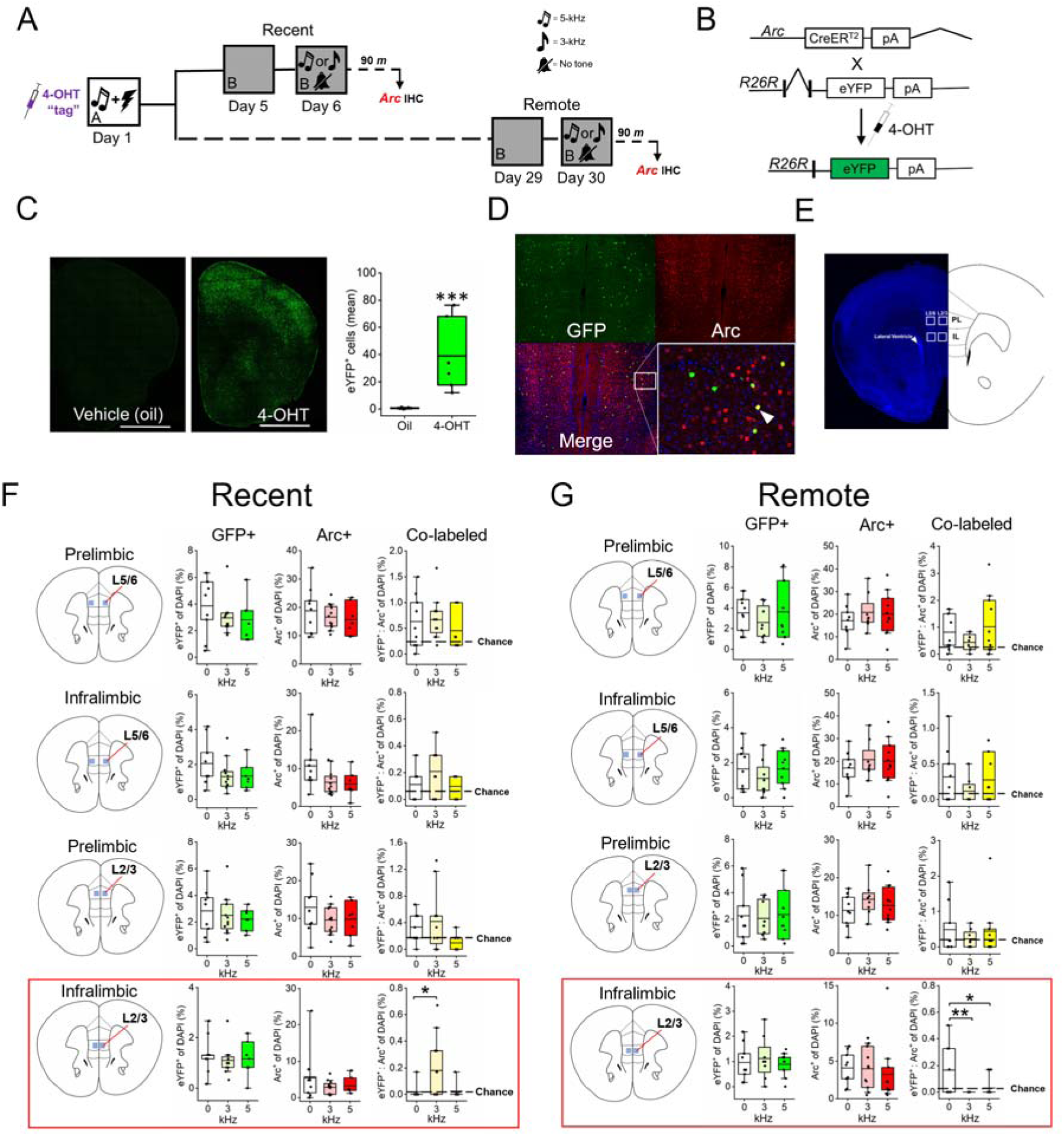
mPFC neuronal ensemble quantitative measures. (A) Experimental design. (B) ArcCreER^T2^ x eYFP system genetic design. (C) Representative photomicrograph depicting the efficacy of 4-OHT to drive eYFP expression after 4-OHT injection versus vehicle (oil) control. (D) 4-OHT (n=6) selectively induces eYFP expression (58-fold increase) versus vehicle (oil; n=6) control in ArcCreER^T2^ mice. (D) Representative photomicrographs depicting GFP and Arc immunofluorescent labeled cells. Arrowhead indicates double-labeled cell. (E) Depiction showing the location of counting frames (250 x 250 microns) in the mPFC. Arrowhead indicates anatomical location of the lateral ventricle reference point. (F) Recent timepoint. Atlas image showing the location of identified neuronal ensemble across PL and IL L2/3 and L5/6. There were no differences in GFP^+^, Arc^+^ or co-localized cells across kHz groups in the PL and IL deep layers. In IL L2/3 (highlighted in red box), there was no difference in eYFP+ or Arc+ expression across groups. However, there was a significant, above chance increase in the number of colocalized cells in the 3-kHz group vs. the no tone control and the CS. (G) Remote timepoint. Atlas image showing the location of identified neuronal ensemble across PL and IL L2/3 and L5/6. There were no differences in GFP^+^, Arc^+^ or co-localized cells across kHz groups in the PL and IL deep layers. In IL L2/3 (highlighted in red box), there was no difference in eYFP+ or Arc+ expression across groups. However, there was a significant, above chance, decrease in the number of colocalized cells in the 3-kHz group and increased above chance number of colocalized cells in the no tone group. n=8-12/group. Hashed lines indicate level of chance calculated as (eYFP +/DAPI+) * (Arc+/DAPI+) * 100. \**p*< .05 and \*\**p*<.01, #*p*<.05 and ##*p*<.01 compared to chance.

To assess whether 4-OHT itself might impact fear learning and memory, a separate group of male and female mice were run through the identical behavioral paradigm described above but instead of 4-OHT, they were injected with vehicle (corn oil) control. These mice were compared with mice injected with 4-OHT. Results showed no differences between Group or Sex interactions during either learning or retrieval of the CS. These data indicate 4-OHT does not impact basic fear memory consolidation behavioral performance.

MANOVA conducted on eYFP^+^ and Arc^+^ cell counts revealed no differences across groups at either timepoint (Figure 3G, H). However, MANOVA conducted on co-activated cells revealed an interaction of Time x kHz *(V*=.49, *F*[4, 40]=4.99; *p*=.002). Follow-up Bonferroni-corrected ANOVAs revealed a kHz x Time interaction in L2/3 IL only (*F*[2,54]=8.0; *p*=.001). At the recent time point (Figure 3F), there was a main effect of kHz (*F*[2,23]=4.7; *p*=.02) (Figure 3G). There were a greater number of co-activated neurons at an above chance level in the GS group versus control (*p*=.04) and no difference between the CS group and controls. Strikingly, a main effect of kHz was also observed at the remote timepoint in L2/3 IL only (*F*[2,25]=6.5; *p*=.005) (Figure 3G). However, in this group, there were fewer co-activated neurons at below chance rates in the GS group versus control (*p*=.004) and also fewer co-active neurons in the CS versus control groups (*p*<.05). Overall, these data indicate that when generalization is low there are more IL L2/3 co-activated neurons but when generalization increases over time there are fewer co-activated neurons in the same location. There were no Time, kHz, or Sex factor differences throughout the PL or IL deep layers at either the recent (Figure 3), or remote (Figure 3), timepoint following learning.

Interestingly, the observed increase in the number of IL L2/3 reactivated cells at an above chance rate in the “no tone” control group at the remote timepoint only (Figure 3H), suggests the possibility that a neuronal ensemble associated with the context elements of the chamber (context fear generalization or extinction) was formed. To address this possibility, we analyzed “context B” habituation data to determine if, 1) mice in the remote group showed increased context fear memory generalization over time, as has been shown previously (Wiltgen & Silva, 2007) and, 2) context extinction retention performance. Analysis of the “context B” behavioral data from the day prior to the cued fear memory generalization test showed a Time x Condition interaction (*F*[3.5, 173.9] = 3.5; *p*=.005). Mice in the remote group showed greater freezing during the first 3 min of the context test relative to mice in the recent group (*F*[1, 52] = 7.6; *p* = .008) (Figure 4A).

**Figure 4.**
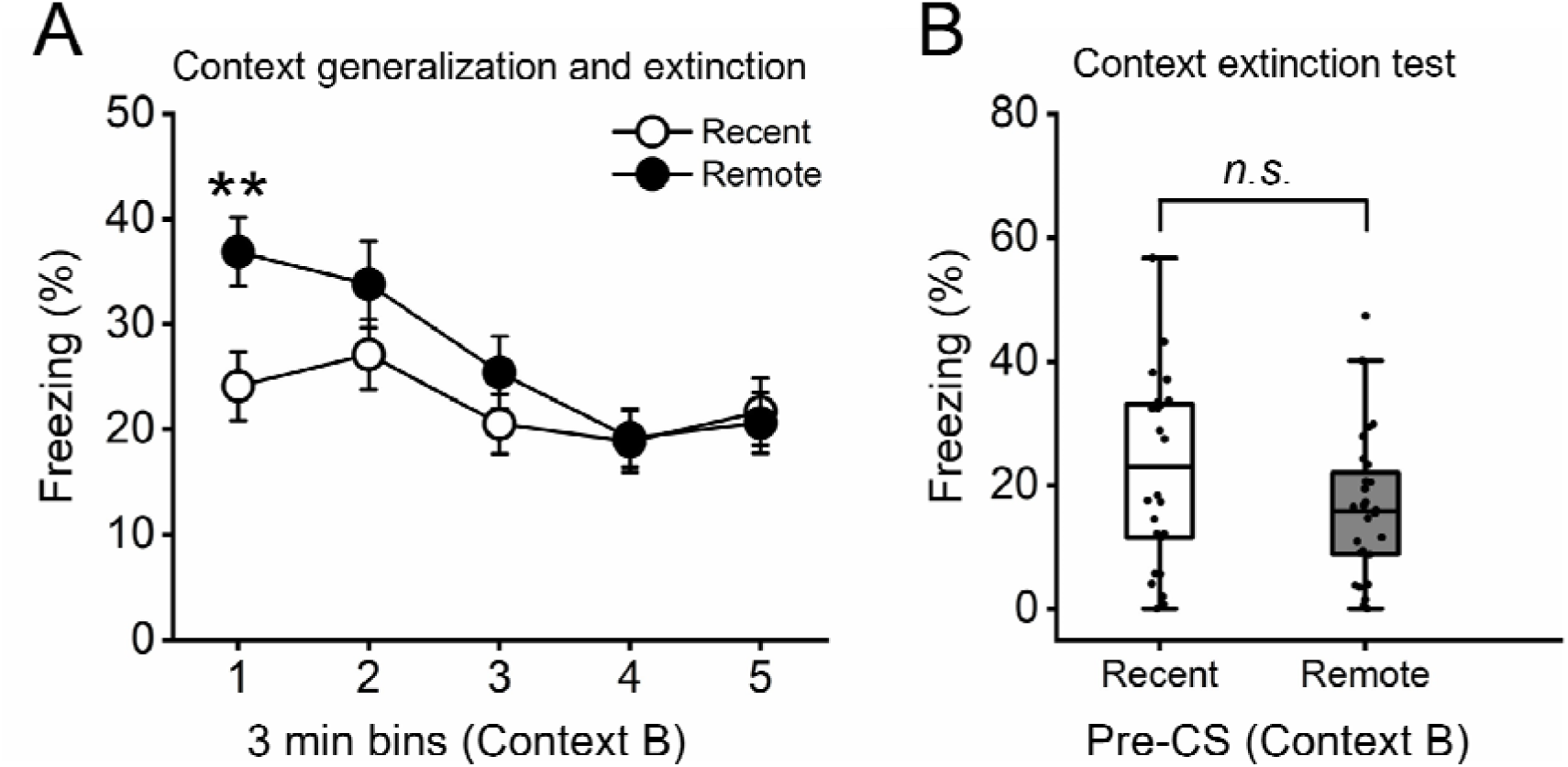
Context generalization and extinction. (A) When were placed into novel context B, mice in the Remote group showed more freezing than the Recent group, indicating context generalization increased with the passage of time. There was also significant extinction over the 15 min session. (B) The next day, during the pre-tone period, there was no differences between groups, indicating strong contextual generalization extinction retention \*\**p*<.01. n=30-31/group

This finding supports previous data showing increased contextual fear memory generalization with the passage of time (Poulos et al., 2016; Wiltgen & Silva, 2007). The difference in the generalized context response declined rapidly over the session and showed no retention the next day (Figure 4B), indicating robust extinction retention of context generalization at the remote time point. This behavioral result was associated with a greater number of reactivated cells during context memory retrieval (pre-GS period), suggesting a neuronal ensemble associated with either greater, 1) context generalization memory or, 2) generalized context fear extinction was formed in IL L2/3 and was expressed during the context test.

### Experiment 4: Bidirectional IL chemogenetic neuronal ensemble control during the expression of cue fear memory generalization

In this experiment, a Cre-dependent DIO-hM4Di, DIO-hM3Dq, or DIO-mCherry control was injected into IL to selectively express DIO-hM4Di, DIO-hM3Dq or DIO-mCherry in ArcCreER^T2^ x eYFP mice (Figure 5A). Subject underwent 4-OHT-dependent tagging during fear conditioning and later retrieval induced neuronal ensemble Arc reactivation was induced using DCZ (Figure 5A).

**Figure 5.**
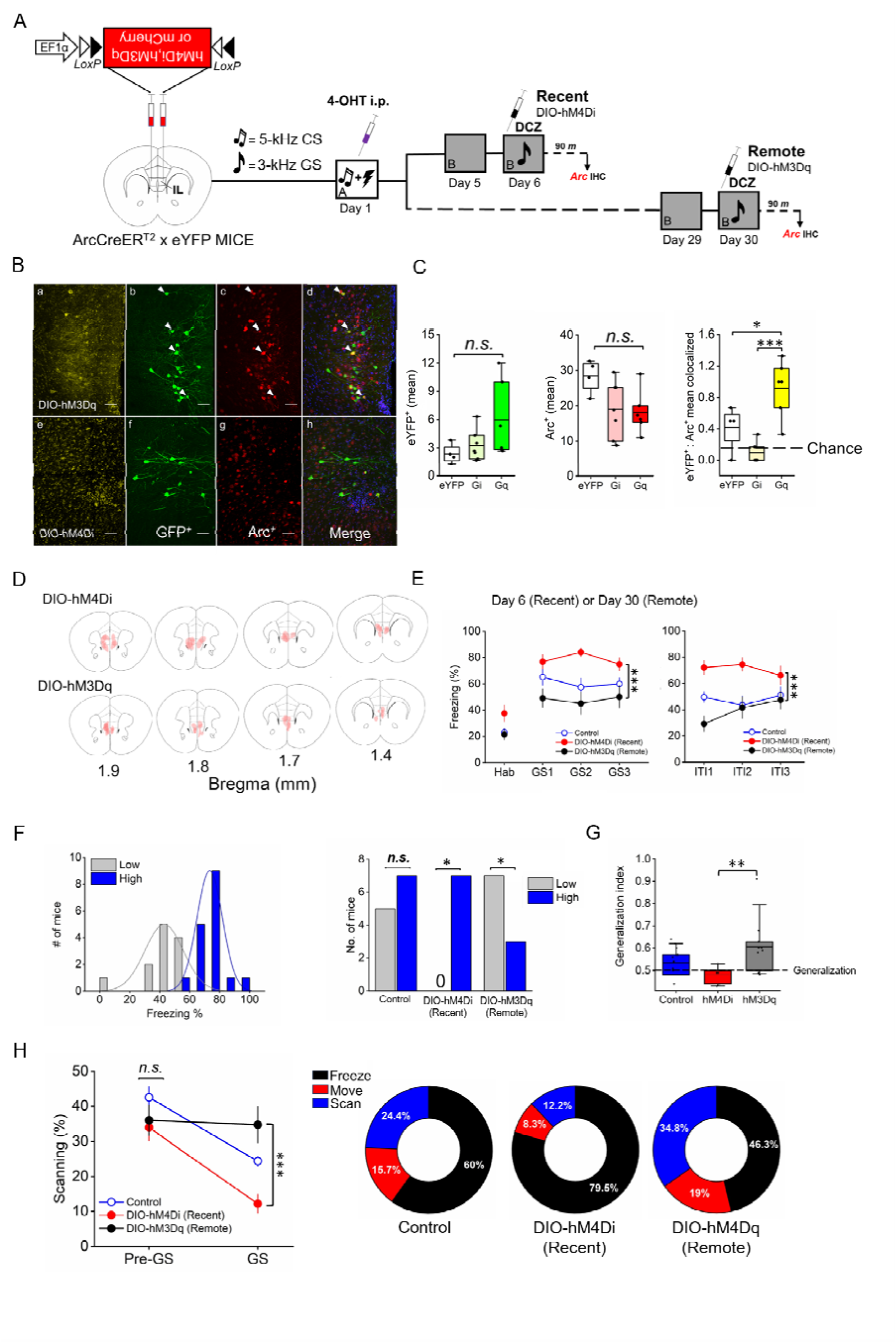
Bidirectional chemogenetic IL neuronal ensemble manipulation during fear generalization expression over time. (A) Schematic depicting DIO-DREADD x ArcCreER^T2^ system design and experimental timeline. (B) Representative photomicrographs showing (a) DIO-hM3Dq AVV transfection in mPFC, (b) GFP^+^-labeled cells, (c) Arc^+^-labeled cells, (d) image merge (20X, 0.8 N.A., 1.5 zoom). Arrowheads indicate double-labeled GFP^+^:Arc^+^ cells. (e) DIO-hM4Di AVV transfection in mPFC, (b) GFP^+^-labeled cells, (c) Arc^+^-labeled cells, (d) image merge. (C) DCZ injection in the DIO-hM3Dq system (n=6) increased the number of co-localized cells relative to DIO-h4Di mice (n=6) and controls (n=4). There was a decrease in DIO-h4Di mice (n=7) versus controls. (D) Heatmapping depicting transfection localized in vmPFC, including IL and DPP. The extent of AAV expression in the mPFC across four coronal planes is depicted. Relatively darker shaded regions show greater overlap across individuals. Transfection was predominantly located in the IL, with some expression in DP. Mouse brain atlas images modified from (Paxinos & Franklin, 2004). (E) We grouped controls as there was no difference in freezing over time and this allowed for a higher-powered cluster analysis. Mice in the DIO-hM3Dq group showed less freezing than DIO-hM4Di mice during presentation of the GS (left panel) and during the ITI (right panel). (F) Unbiased cluster analysis on the mean GS freezing response across all mice produced 2 prominent clusters (Low and High). Chi-square analysis indicated all mice in the DIO-hM4Di were clustered in the “High” group and a majority of mice in the DIO-hM3Dq groups clustered in the “Low” group. (G) Analysis of the generalization index confirmed these results. (H) Prior to GS presentation, there was no differences in scanning behavior between groups. Upon presentation of the GS, scanning collapsed in the DIO-hM4Di group but was maintained in the DIO-hM3Dq group. Donut plots depict percentage freezing, scanning, and moving during GS presentation (right panel). Mice in the DIO-hM3Dq spent more time scanning and moving compared with DIO-hM4Di mice. n=7-14/group. \**p*<.05, \*\**p*<.01, \*\*\**p*<.001. Scale bar = 50 *µ*m.

The efficacy of DCZ to modulate neuronal activity in DIO-DREADD expressing cells, was assayed using IHC against eYFP and Arc. Results revealed no differences in eYFP^+^ or Arc^+^ neuronal expression across groups (Figure 5B-C). When the number of eYFP^+^:Arc^+^ co-localized cells was analyzed, we found a difference across groups (*F*[2, 14]=15.2; *p*<.001). There were a greater number of co-activated cells in the DIO-hM3Dq group compared with the control (*p*<.05) and DIO-hM4Di group (*p*<.001) (Figure 5C). These data indicate the combination of DCZ and the DIO-DREADD system preferentially drives Arc^+^ expression in eYFP-labeled cells. Analysis plotting the spread of viral transfection across brains showed clear AAV transfection in IL and also in DP (Figure 5D). We grouped both bilateral and unilateral transfection into the analysis because there was a high degree of laterality in the expression of eYFP which may have anatomically biased transfection. In addition, there were no statistical differences between bilateral and unilateral were detected between groups. We also grouped controls as there was no difference in freezing over time and this allowed for a higher-powered cluster analysis.

By chemogenetically manipulating the activity of a neuronal ensemble established during learning, here we test its contribution to the expression of generalization. Results revealed a main effect of DIO-DREADD manipulation (*F*[3, 28]=12.0; *p*<.001). Mice in the DIO-hM3Dq remote group showed less GS-elicited freezing than mice in the DIO-hM4Di recent group (*p*=.002) (Figure 5E). Like the previous global DREADDs analysis, an unbiased k-means clustering algorithm was applied to mean GS freezing levels across all mice. The analysis revealed 2 prominent “High and Low” clusters (Figure 5F). A chi-square test was performed to test the relationship between experimental groups and clusters. Results showed a significant effect ^2^ (2, N=29) = 8.3, *p*=.016, with mice in the DIO-hM4Di group solely clustered in the “High” generalization group while mice in the DIO-hM3Dq group clustered predominately in the “Low (n=7)” with some mice clustering in the “High” generalization group (n=3). Analysis of the generalization index revealed a significant difference across group (*F*[2, 28]=5.6; *p*=.009), with mice in the DIO-hM3q groups exhibiting greater scores (less generalization) than mice in the DIO-hM4Di group (Figure 5G). For scanning behavior, RMANOVA showed a Time x DIO-DREADD interaction (F[2, 35]=3.7; *p*=.036). There were no differences in scanning behavior prior to the presentation of the GS across all groups (Figure 1H). However, upon GS presentation, scanning behavior collapsed in the DIO-hM4Di and control group, but was maintained in the DIO-hM3Dq (*F*[4, 32]=9.04; *p*<.001) (Figure 1H). This result replicates findings also seen in Experiment 1. Overall, these results indicate neuronal ensembles established in the IL during learning contribute to later conditioned generalization. However, given DP transfection in a few of our subjects (Figure 5D), interpretation of these results cannot rule out DP functionality in generalization. Throughout the experiments, no sex interactions were detected.

## DISCUSSION

Here we leveraged an ArcCreER^T2^ x eYFP transgenic system to genetically tag IL neurons activated during Pavlovian fear conditioning for later visualization and manipulation during the expression of generalization. Shortly after learning, more reactivated IL neurons were associated with the GS and silencing an IL neuronal ensemble resulted in more generalization and a collapse in scanning behavior. Conversely, later after learning, fewer reactivated IL neurons were associated with the GS, and stimulating an IL neuronal ensemble promoted less generalized freezing and more scanning. One interpretation of these results is that, at the recent timepoint following learning, presentation of a similar, but novel, GS promoted IL neuronal ensemble reactivation and memory specificity, presumably because memory for the original CS was intact. Over time, some attribute of the CS (i.e., frequency) may have been forgotten (Bouton et al., 1999), resulting in reactivation failure and the expression of generalization. A key outstanding question is whether the identical IL neuronal ensemble reactivated in response to the GS presented shortly after learning was suppressed at the remote timepoint after learning.

An intriguing hypothesis of these data is that an IL neuronal ensemble encoding an inhibitory process for attenuating generalized freezing was formed in the IL during initial memory consolidation and expressed in the presence of the ambiguous GS. When this ensemble failed to reactivate, generalization increased. This empirical finding confirms a long-standing theory proposing generalization as a learning process (Hull, 1943). There is also recent work showing IL stimulation (picrotoxin) after learning improved later extinction performance and protein synthesis inhibition eliminated the effect. These data suggest that memory consolidation for extinction is established following learning (Bayer et al., 2024). We theorize that a similar “opponent process,” as that described by Bayer et al. (2024), may have been formed in the IL during learning to later modulate generalization.

Overall, these data support previous findings showing, 1) that the passage of time increases generalization (Pollack et al., 2018; Poulos et al., 2016; Wiltgen & Silva, 2007), 2) a selective role for the IL in processing uncertain threat stimuli (Glover et al., 2020) and, 3) involvement of IL L2/3 plasticity in generalization processes (Pollack et al., 2018; Scarlata et al., 2019).

### Cued fear memory expression, cued extinction, generalized contextual extinction, and IL functionality

Cued fear memory expression was spared during IL stimulation. However, on the next day without CNO, cued fear memory expression decreased. While this effect was small, we speculate that a brief CS exposure schedule (3 random ITI presentations) may have acted as a “weak” extinction training regimen that, in combination with IL chemogenetic stimulation, promoted stronger extinction retention the next day. This finding supports IL functionality in fear extinction consolidation rather than learning (Bayer & Bertoglio, 2020; Bloodgood et al., 2018; Bukalo et al., 2015) and another study showing that increasing IL activity (using picrotoxin) normalized extinction retrieval in an extinction-impaired mouse strain (Fitzgerald et al., 2014). Interestingly, we did not find increased ensemble reactivation in response to the CS at either recent or remote timepoints. This finding may be supported by data showing the IL is not tone responsive during extinction training (Milad & Quirk, 2002).

At the remote timepoint only, the “no tone” control group was associated with more L2/3 reactivated cells. We hypothesize this ensemble was associated with the emergence of a new contextual fear extinction memory. The reasoning behind this hypothesis is that all mice were placed into alternate “context B” a day prior to memory reactivation which effectively served as a context generalization test and extinction session. Analysis of these data showed increased context generalization over time, a finding that supports previous work (Wiltgen & Silva, 2007). There was also a reduction in freezing over the test session (extinction) and no difference in freezing on the generalization test day over time between recent and remote groups, indicating “strong” contextual fear memory extinction retention. Together, these data suggest a separate IL L2/3 neuronal ensemble was formed and associated with context generalization extinction. Further, fewer co-activated cells in both GS and CS groups implies that the presentation of the tone stimuli may have suppressed reactivation related to contextual extinction. Overall, these data support a role for the IL in context fear extinction (Brockway et al., 2023), context generalization (Bayer & Bertoglio, 2020), and the extinction of generalized contextual fear.

### Memory precision and a theoretical basis for dynamic changes in IL ensemble activity over time

At the remote timepoint following learning, neuronal ensemble activity in response to the GS was reduced. There are several plasticity-related mechanistic possibilities that might account for this phenomenon: 1) IL gradually disengaged its obligatory top-down inhibitory role in the defensive memory circuit over time to support increased generalization, in a systems consolidation-like process (Bergstrom, 2016), 2) IL activity or IL neuronal ensemble reactivation may have been suppressed over time via newly recruited inhibitory mechanisms (Morrison et al., 2016), or 3) IL neuronal ensemble activity may have weakened over time through a synapse destabilization mechanism, in a process akin to forgetting (Ryan & Frankland, 2022).

With respect to possibilities 2 and 3, one proposal is that neuronal networks employing mechanisms of instability, such as synaptic decay, elimination, or suppression, would promote a greater degree of generalization (Richards & Frankland, 2017). A weakening of synapses (destabilization) may provide a “forgetting” mechanism to promote brain states that reflect the past. That is, in the face of threat ambiguity and over time, it may be advantageous to promote or bias the original CR to enhance the chance of survival in “a better safe than sorry” strategy (Eilam et al., 2011). The exact nature of the suppression of IL L2/3 neuronal ensemble reactivation with respect to generalization processes remains an open question.

### Risk assessment, switching defensive states, and IL functionality

In the presence of potential threat, organisms must first detect the threat and then respond, with the most appropriate defensive stance. Predatory Imminence Continuum Theory organizes potential defense responses along a spectrum, from safety (i.e., movement related to foraging), to pre-encounter threat (i.e., vigilant behavior such as scanning), to post-encounter threat (i.e., freezing), to threat contact (i.e., circa-strike defense movement related to fight, flight, or panic) (Fanselow, 1994). Here, we hypothesized that mice presented with the GS and exhibiting low levels of freezing might engage alternate defensive responses based on the relative ambiguity of perceived threat. Vigilant scanning behavior was identified as a distinguishable, and highly prominent, defensive behavior that aligns with a pre-encounter threat response to the ambiguous tone (Blanchard et al., 2011). When IL was stimulated, scanning behavior made up a substantial portion (over 34%) of all movement. Conversely, freezing (generalization) was the predominant response when the IL was inhibited. These data indicate IL functionality, and IL neuronal ensembles, in switching defensive states between post- and pre-encounter; IL inhibition resulted in post-encounter, and IL stimulation promoted pre-encounter, defensive states in response to the GS, potentially via downstream signaling with central nucleus of the amygdala (Moscarello & Penzo, 2022). These findings support other work indicating IL in switching behavioral defensive states to promote movement (Halladay & Blair, 2016) and also theoretical work suggesting a role for IL in promoting behaviors that align with the most rigid reading of the associative relationship amongst cues present (Nett & LaLumiere, 2021). While freezing and scanning are categorized as risk assessment behaviors, they are differentiated by the actual or perceived proximity or ambiguity of the threat.

### Clinical relevance

The use of novel “ambiguous” stimuli in an auditory cued Pavlovian defensive conditioning paradigm to study brain functionality underlying generalization processes has preclinical utility for the study of PTSD (Dunsmoor & Paz, 2015) and anxiety disorders (Lissek et al., 2006). In the field of psychology, a “weak situation” refers to a context in which environmental stimuli are less predictive of threat imminence. The ambiguous nature of the environment tends to produce more individual difference variability and has been proposed as a model for studying anxiety (Lissek et al., 2006). In our paradigm, both the contextual elements and auditory tone frequency were shifted, which might account for wide variation in individual differences. Considering the negative valence domain within the Research Domain Criteria (RDoC) framework, the study of potential, but ambiguous, threat has implications for the study of anxiety that can be differentiated from acute threat (Fanselow & Hoffman, 2024). The vmPFC functions in the expression of generalization in humans (Spalding, 2017) and vmPFC dysfunction has been associated with PTSD and anxiety-related disorders (Alexandra Kredlow et al., 2022). The consensus of data presented here indicates vmPFC functionality, and perhaps vmPFC neuronal ensembles, in “down-shifting” defensive behavioral states. IL is considered a putative analog of Area 25 in humans, which regulates mood and emotion (Alexander et al., 2019). Thus, the methodology and data in this set of experiments may add preclinical value for understanding the role of IL (Area 25) dysfunctionality and over-generalization phenomenon seen in anxiety and trauma-related disorders.

## Acknowledgments

Authors thank Dr. Lindsay Halladay for reviewing a previous version of the manuscript. Authors thank Kate Carter for experimental technical expertise, Dave Lewis for microscopy expertise, and the Vassar College Animals Resources team, including Paul Gonzalez, Yi Yaun, Gina Coluccio, and Leah Erickson, for expert care of animal resources.

