## Supplementa material for "An infralimbic cortex neuronal ensemble encoded during learning attenuates fear generalization expression" (from another version of this preprint: version 2, 2024-08-20)

### Supplemental figures

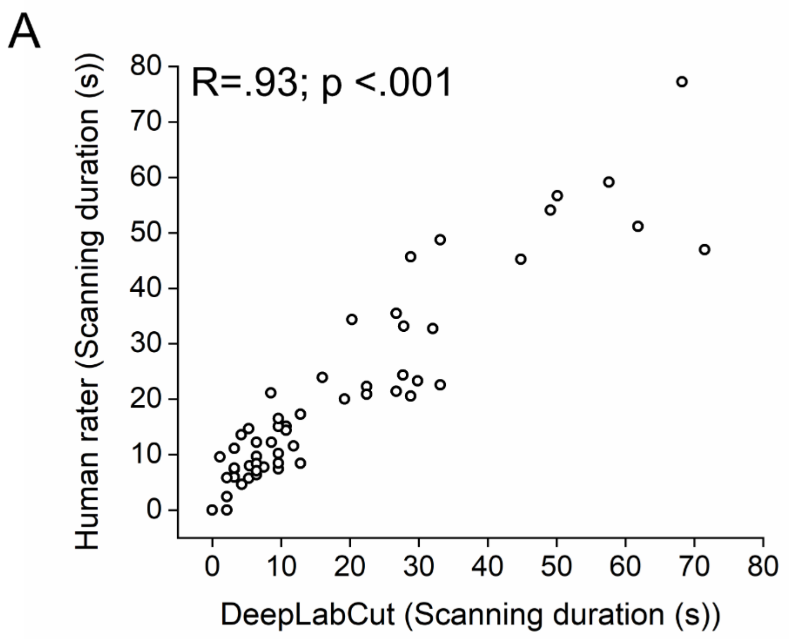


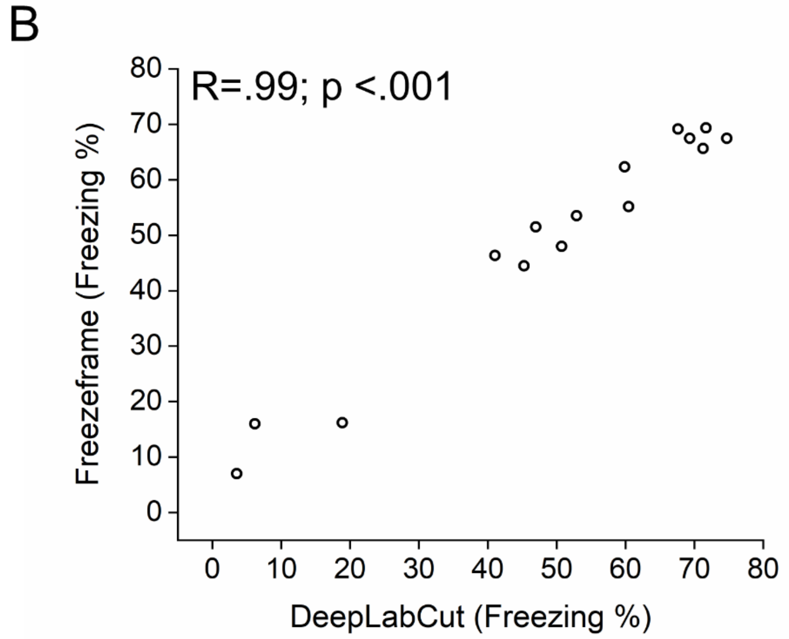


**Figure S1**. Inter-rater reliability measures (Pearson's correlation). Results showed high correlation between DLC scanning analysis and a human rater, n=56 for human vs. DLC (A) and DLC freezing analysis compared with Freezeframe, n=21 for Freezeframe vs. DLC. (Actimetrics, Lafayette, ID)

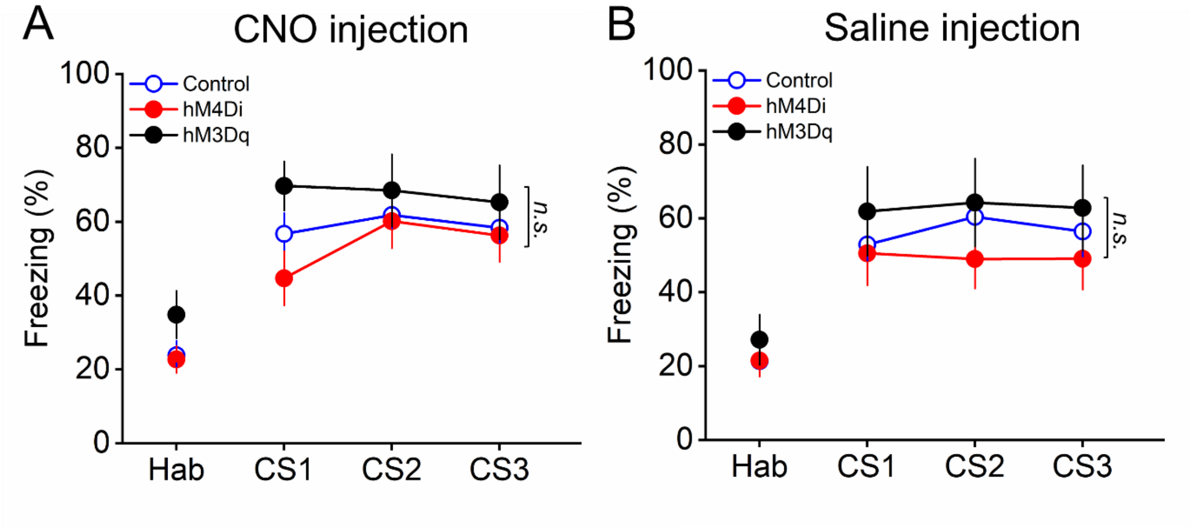


**Figure S2.** There were no significant differences in generalized freezing in mice with unilateral DREADD transfection and injected with CNO (A) and saline (B) across hM4Di (n=17), hM3Dq (n=10), and control (n=27) groups. n.s. non-significant statistical difference.

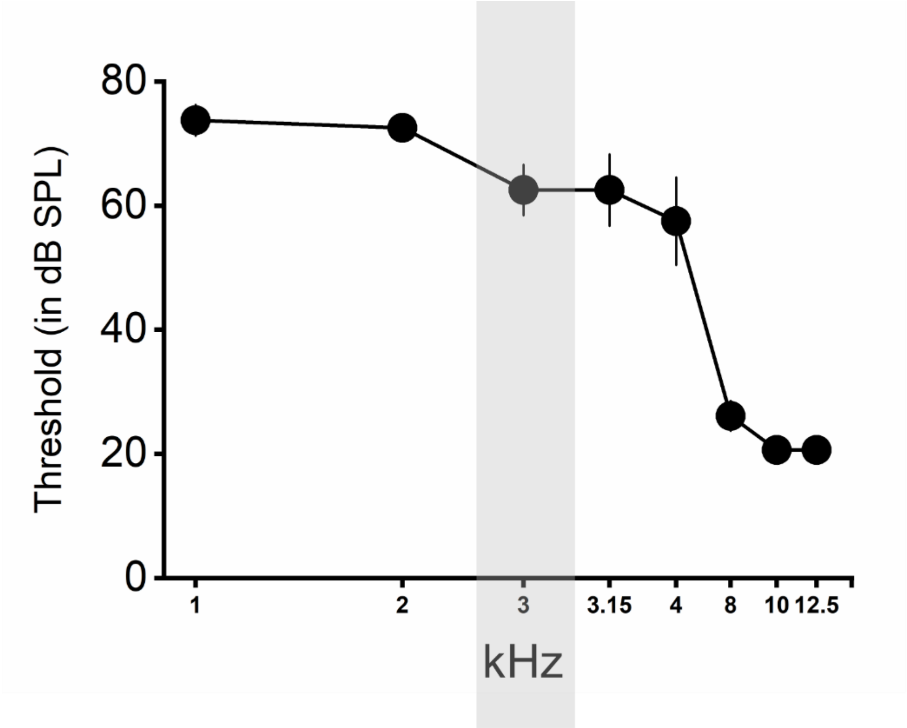


**Figure S3:** Brainstem auditory evoked potentials. Thresholds at 3 kHz ranged from 57.5 to 67.5 dB SPL, with an average of 62.5 dB SPL, which are lower than the amplitude of the stimulus used in the generalization experiments (70-75 dB SPL). n=4

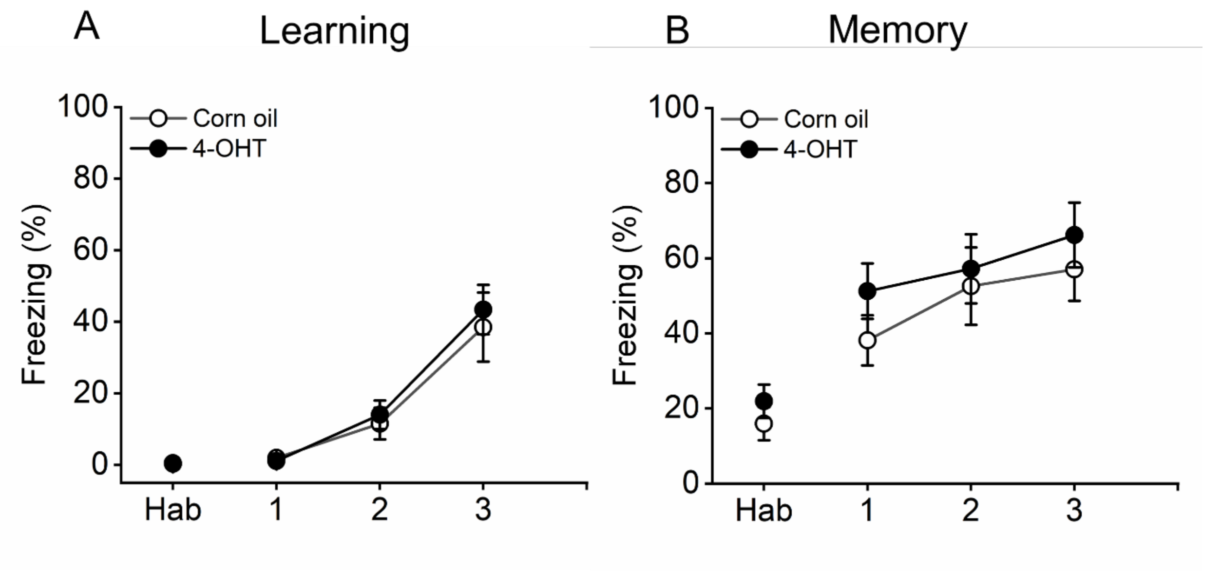


**Figure S4.** Mice injected with 4-OHT showed similar learning (A) and memory (B) performance compared with a corn oil vehicle control injection. (oil; n = 12; female = 7 and male n = 5). These mice were compared with mice injected with 4-OHT (n = 13; female n = 7 and male n = 6).

### Recent

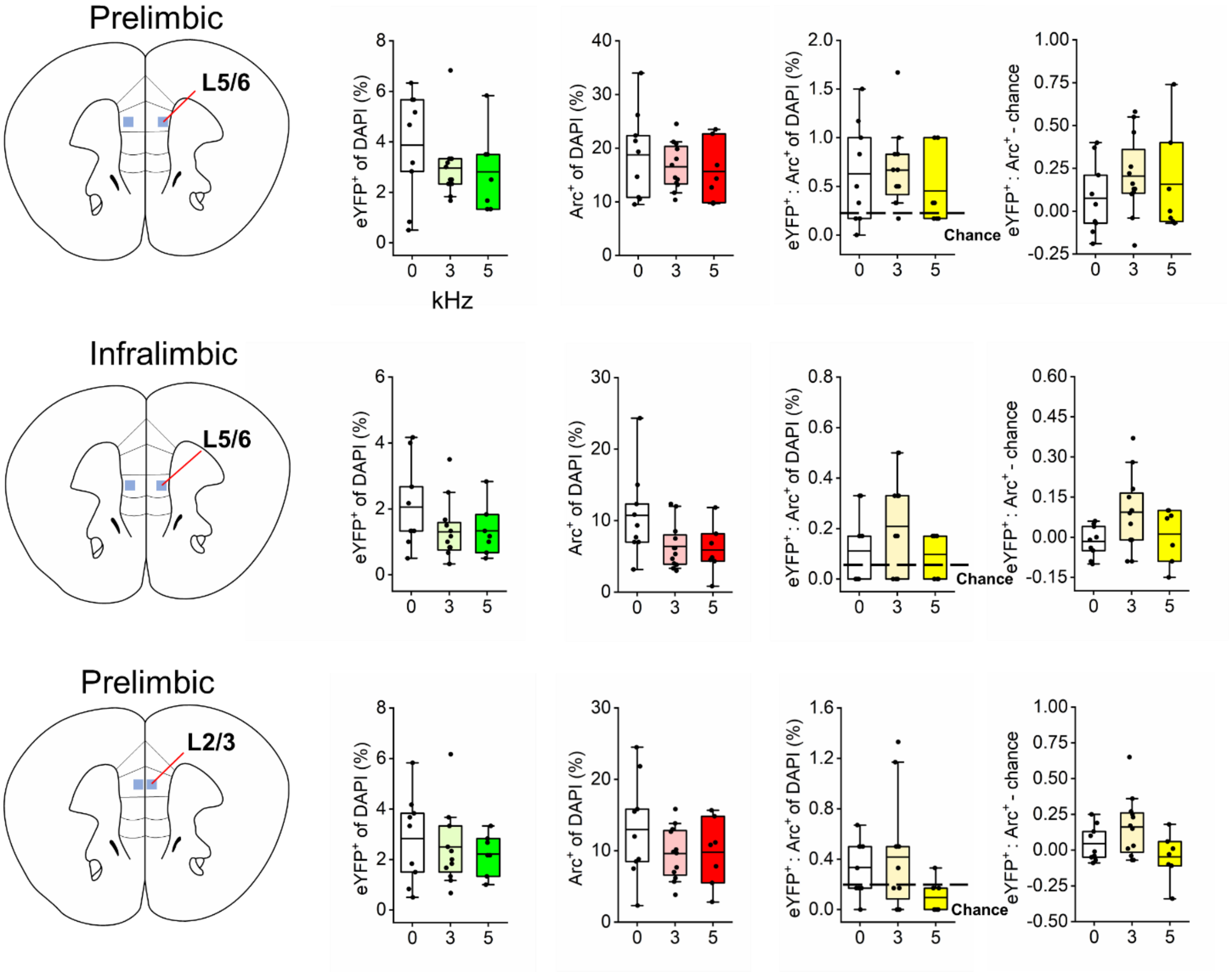


**Figure S5.** At the recent timepoint, there was no difference in GFP+ or Arc+ expression across groups in both PL deep and shallow layers and IL deep layers. There was a significant, above chance decrease in the number of colocalized cells in the 3-kHz group and increased above chance number of colocalized cells in the no tone group in IL L2/3 #p<.05 compared to chance. **p<.01 and *p<.05 compared with no tone

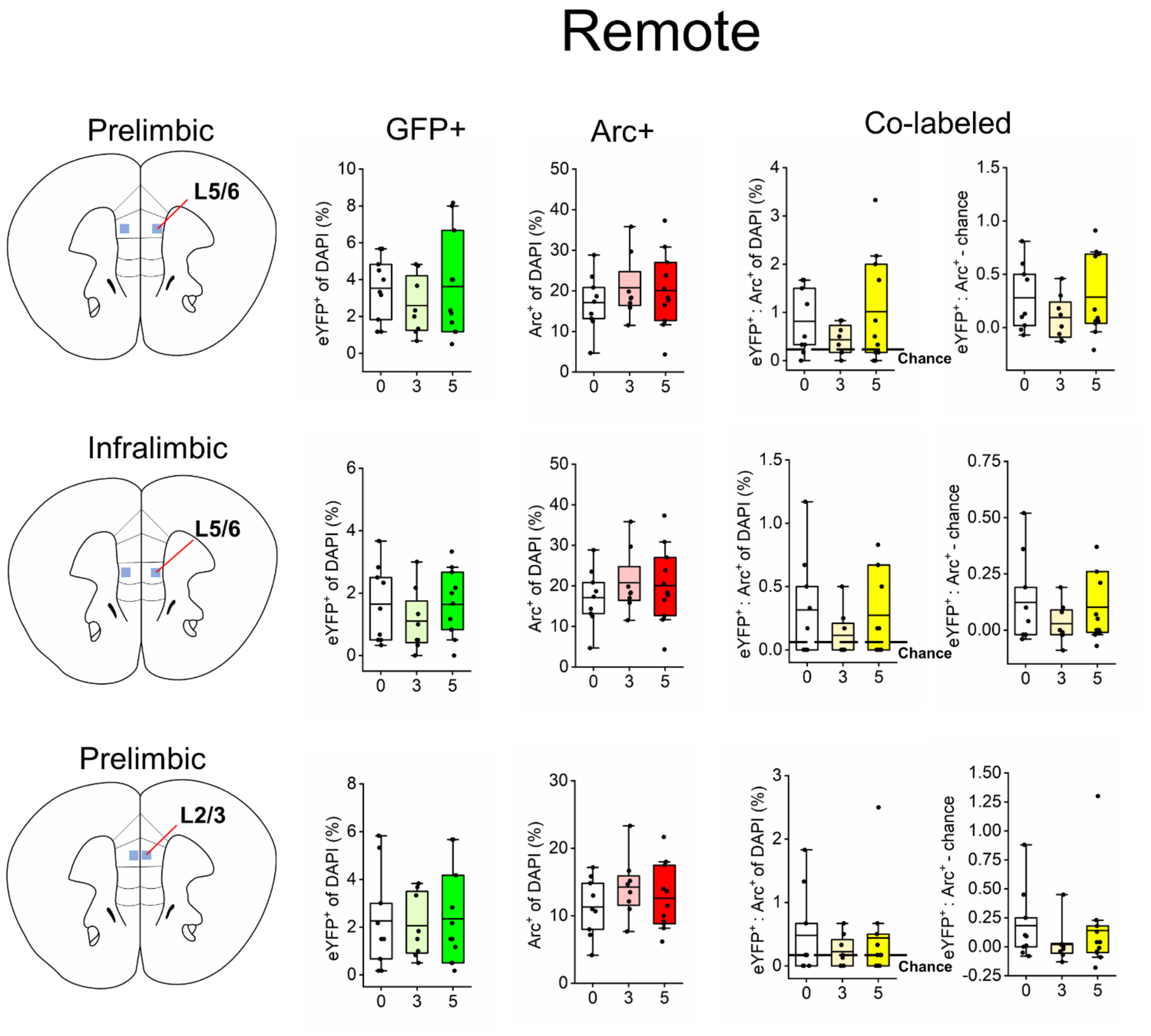


**Figure S6.** At the remote timepoint, there was no difference in GFP+ or Arc+ expression across groups in both PL deep and shallow layers and IL deep layers. There was a significant, above chance decrease in the number of colocalized cells in the 3-kHz group and increased above chance number of colocalized cells in the no tone group in IL L2/3 #p<.05 compared to chance. **p<.01 and *p<.05 compared with no tone
